# A structural Merton jump-diffusion framework for survival analysis: Modeling biological solvency and distance-to-death(DtD) in tuberculosis

**DOI:** 10.64898/2026.03.30.715204

**Authors:** Eric Walter Pefura-Yone, Eric Herman Pefura-Yone, Hillary Laeticia Njonap Pefura-Yone, Amadou Djenabou, Adamou Dodo Balkissou

## Abstract

Tuberculosis (TB) remains a leading cause of death globally, with early mortality often driven by severe malnutrition and human immuno-deficiency virus (HIV) co-infection. Traditional survival analyses identify risk factors but remain associative, failing to capture the dynamic physiological collapse preceding death. In a novel interdisciplinary adaptation, we applied the Merton jump-diffusion structural framework from quantitative finance to model survival as a state of biological solvency, in which mortality occurs when a stochastic health trajectory crosses a critical failure threshold. We analysed a retrospective cohort of 15,182 TB patients in Cameroon over two decades. Adjusted body mass index (BMI) was conceptualized as a proxy for health capital and modeled using a stochastic process accounting for individual recovery trends, physiological instability, and acute clinical shocks. The study included predominantly young adult males (median age: 33 years) with a median BMI of 20.7 kg/m^2^. HIV co-infection was present in 35% of patients. The overall mortality rate during the 240 days follow-up period was 7.0%, with 55.1% of deaths occurring within the first 30 days. The model identified a critical failure threshold at BMI 17.329 kg/m^2^. HIV co-infection emerged as a key driver of metabolic instability, significantly increasing physiological volatility. Statistical validation confirmed that sudden clinical shocks were necessary to explain observed mortality patterns. The resulting Distance-to-Death (DtD) metric slightly outperformed standard associative extended Cox models in predicting survival, achieving a higher discriminative ability in testing set (Harrell’s C-index: 0.781 vs. 0.772; p = 0.038). Patients stratified into the highest-risk category showed a mortality rate of 16.7%, compared with 1.6% in the most stable group.This study bridges financial engineering and clinical epidemiology, offering a mechanistic understanding of how physiological reserves and metabolic instability determine survival. To support clinical application, we developed an interactive digital triage tool enabling identification of high-risk patients in resource-limited settings.

**Author summary:** Tuberculosis remains a major cause of death worldwide, particularly in people with poor nutrition or co-infection with HIV. In this study, we explored a new way to understand why some patients survive while others do not. We adapted a method originally used in finance to track the “health reserves” of patients over time, using body weight and related measures to estimate how close someone is to a critical health threshold. Our approach captures both gradual health decline and sudden medical complications, such as severe infections or rapid deterioration. By applying this method to a large group of patients in Cameroon, we found that a very low body weight is a strong warning sign for impending death and that HIV infection makes health outcomes less predictable. We also created a simple scoring tool that can help doctors identify patients at greatest risk, so that life-saving interventions and closer monitoring can be prioritized. This work bridges mathematical modeling and clinical care, offering a new way to assess patient vulnerability and improve outcomes in resource-limited settings.

## 1. Introduction

Tuberculosis (TB) remains one of the leading causes of infectious mortality worldwide, with a disproportionate burden in resource-limited settings[1]. Despite therapeutic advances, early mortality remains high, particularly among patients presenting with severe malnutrition, HIV co-infection, or complicated clinical forms[2–5]. Consequently, the early identification of patients at high risk of death remains a major public health challenge. Among the clinical markers available at admission, the body mass index (BMI) occupies a central role. Numerous studies have shown that a low BMI is strongly associated with an increased risk of death in TB patients, independent of other clinical and sociodemographic factors[6,7]. BMI reflects nutritional status, physiological reserve, and the systemic impact of the infection, making it a particularly relevant integrative biomarker in this context[6–11].

Traditionally, the risk of death in TB patients is analysed using semi-parametric survival models, most notably the Cox proportional hazards model[12]. These approaches have significantly contributed to the identification of factors associated with mortality and remain the gold standard in clinical epidemiology. However, even when extended to incorporate time-varying covariates or non-proportional hazards, Cox-based survival models remain fundamentally associative, as they estimate the marginal effects of covariates on the instantaneous hazard without explicitly modeling the underlying temporal dynamics of continuous biomarkers or the pathophysiological processes leading to death[13].

In parallel, other disciplines facing failure-related problems, notably quantitative finance and reliability engineering, have developed structural models based on continuous stochastic processes. The Merton model, initially proposed to quantify corporate default risk, posits that a system evolves randomly around an average trend and that a critical event occurs when this process crosses a predefined barrier[14]. This framework introduced the concept of “Distance to Default” (DtD), which has become a central tool for risk assessment. The KMV model further operationalizes the Merton framework by calibrating it to observed data, producing actionable default probabilities for individual firms[15,16]. Analogously, structural frameworks, such as the Merton Jump-Diffusion model, can be employed in the analysis of clinical data, thereby modeling the stochastic progression of physiological reserves across temporal dimensions. These models effectively encapsulate both progressive decline and sudden disruptions, thereby furnishing a mechanistic understanding of individual mortality risk that goes beyond traditional associative Cox models.

Conceptually, this type of modeling presents striking analogies with certain medical situations. In the case of TB, BMI can be interpreted as a dynamic physiological reserve, subject to recovery or deterioration trends, individual variability (metabolic instability), and occasional disruptions from acute clinical shocks such as hospitalization or severe complications. Death can thus be envisioned as the crossing of a critical threshold of physiological fragility. To date, the explicit use of structural models inspired by finance to model the risk of death in epidemiology remains very limited. Aalen and Gjessing (2001) highlighted that modeling survival as a stochastic trajectory, incorporating concepts such as quasi-stationary distributions in Markov or diffusion processes, provides deeper insight into individual risk dynamics[17]. Such approaches have been largely underutilized in biomedical research, yet they offer a mechanistic perspective on mortality that goes beyond traditional associative models.

Building on this conceptual foundation, our work operationalizes the idea by applying a Merton Jump-Diffusion framework. We present a proof-of-concept that transposes Merton’s structural model to clinical epidemiology by modeling the risk of death in TB patients. By treating the BMI trajectory as a continuous stochastic process with drift, volatility, and jumps, we introduce the concept of “Distance to Death” (DtD), analogous to the “Distance to Default” in finance. The objective is not to replace existing survival models, but to demonstrate the mathematical coherence, clinical interpretability, and epidemiological relevance of this approach in an exploratory framework. This study aims to open a new methodological perspective on risk assessment in infectious diseases, proposing a conceptual bridge between structural modeling and clinical epidemiology. The primary goal was to model mortality risk during the intensive and continuation phases of TB treatment using a structural stochastic framework.

## 2. Materials and Methods

### 2.1. Study design and participants

We conducted a large-scale retrospective cohort study at Jamot Hospital of Yaoundé (YJH), the national reference center for respiratory diseases and TB care in Cameroon. The study was designed to capture long-term treatment trajectories and survival dynamics of patients undergoing anti-tuberculosis therapy in a high-burden setting. The Diagnostic and Treatment Center (DTC) for TB of HJY functions as a major referral hub within the National Tuberculosis Control Program (NTCP), managing approximately 1,500 to 1,800 TB cases annually. This high patient throughput, combined with standardized treatment and follow-up procedures, provides a robust empirical foundation for modeling complex time-to-event processes, including treatment failure, loss to follow-up, and mortality. All patients with TB aged ≥ 15 years treated with World Health Organisation (WHO) standard regimens for non rifampicin or non multidrug resistant TB [18] between January 2001 and December 2021(20 years) were included in this study. Patients with missing data on key variables (clinical form of TB, status for HIV infection, body weight) were excluded. Ethical clearance was granted by the Faculty of Medicine and Biomedical Sciences, University of Yaoundé I (Approval No. 0195/2023).

### 2.2. Classification of TB cases

At YJH, the diagnosis and classification of TB follow the recommendations of the World Health Organization (WHO) and the Cameroon National Tuberculosis Control Program (NTCP)[19,20]. Standardized international definitions are applied at the DTC to ensure consistency in case identification and clinical management. Pulmonary TB is classified as smear-positive (SPPTB) when acid-fast bacilli (AFB) are detected in at least one sputum specimen. Smear-negative pulmonary tuberculosis (SNPTB) is defined by persistently negative sputum smear examinations after a 10-day course of non specific antibiotic therapy in patients presenting with clinical and radiological features suggestive of tuberculosis, in the absence of an alternative diagnosis, and based on the clinician’s decision to initiate a full course of anti-tuberculosis treatment. Extra-pulmonary tuberculosis (EPTB) refers to tuberculosis involving organs other than the lungs. Patients with a history of prior anti-tuberculosis treatment are classified as retreatment cases and further categorized as relapse (recurrence of disease after successful completion of a previous treatment course), treatment failure (persistence of smear positivity at or beyond five months of therapy), or treatment after loss to follow-up (resumption of treatment after an interruption of at least two consecutive months). A new case is defined as a patient who has never received anti-tuberculosis treatment or who has been treated for less than one month. Other cases of tuberculosis include patients who do not meet the criteria for any of the aforementioned categories. In accordance with NTCP guidelines, all retreatment cases managed at this centre are systematically screened for multidrug-resistant tuberculosis using either phenotypic drug susceptibility testing or the GeneXpert MTB/RIF assay.

### 2.3. TB treatment and outcomes

Management of active tuberculosis followed the WHO and national guidelines issued by the Cameroon NTCP and reflected the standard-of-care practices in place during the study period[18–20]. Depending on disease severity and patient circumstances, treatment was delivered either on an inpatient or outpatient basis during the intensive phase. Newly diagnosed patients with drug-susceptible tuberculosis received the standard first-line regimen over six months, consisting of a two-month intensive phase with rifampicin (R), isoniazid (H), ethambutol (E), and pyrazinamide (Z), followed by a four-month continuation phase with rifampicin and isoniazid (2RHEZ/4RH). During the early phase of the study, retreatment cases were managed using an eight-month regimen that included R, H, E, and Z, supplemented with streptomycin (S) during the first two months and pyrazinamide during the first three months (2RHEZS/1RHEZ/5RHE). Between 2017 and 2018, in response to evolving WHO guidance, streptomycin-containing regimens were discontinued. Retreatment cases without confirmed rifampicin resistance or multidrug resistance were thereafter treated using the standard six-month first-line regimen (2RHEZ/4RH). From 2019 onward, retreatment patients with non rifampicin resistance received the simplified six-month regimen (6RHEZ). In accordance with national policy, all tuberculosis patients were offered HIV testing after obtaining informed consent. HIV-positive individuals received cotrimoxazole prophylaxis and were initiated on combination antiretroviral therapy (cART) free of charge, typically within two weeks of diagnosis, consistent with integrated TB/HIV care guidelines.

At the end of follow-up, treatment outcomes were classified into mutually exclusive categories following NTCP and WHO definitions[18,20]. Patients were considered cured if they had a documented negative sputum smear at the end of treatment and at least one previous negative result during therapy. Treatment completion applied to patients who completed the prescribed regimen without bacteriological confirmation at the final visit. Treatment failure was defined as persistent or recurrent smear positivity from the fifth month onward. Death included mortality from any cause during treatment. Loss to follow-up was defined as interruption of therapy for two or more consecutive months, and transfer-out referred to patients referred to another treatment centre with unknown final outcomes. For analytical purposes, cured patients and those who completed treatment were considered to have achieved a favourable outcome.

### 2.4. Data collection

Data were obtained through a review of anti-tuberculosis treatment registers and individual patient charts. Collected information included sociodemographic characteristics (age and sex); clinical characteristics such as initial hospitalization status, tuberculosis form (smear-positive pulmonary TB, smear-negative pulmonary TB, or extrapulmonary TB), patient category (new or retreatment case), and HIV serostatus; and anthropometric measures, including body weight and an adjusted BMI, calculated as weight divided by the square of the mean adult height by sex (1.70 m for men, 1.60 m for women). This BMI adjustment accounts for sex differences in body composition while preserving its validity as a proxy for physiological reserve[21]. Treatment outcomes (success, death, failure, or loss to follow-up) and duration of follow-up were also recorded.

### 2.5. Conceptual framework of the Merton model

To establish a robust conceptual model linking TB clinical variables to financial risk modeling, patient health is conceptualized as a “firm”, and survival is interpreted as biological “solvency”. Within this conceptual framework, we apply the Merton jump-diffusion structural model, originally developed to predict corporate default[14,16]. Clinical default is defined as death occurring during anti-tuberculosis treatment. This approach enables a mechanistic interpretation of disease progression by translating physiological deterioration into a formal solvency-based risk structure.

#### 2.5.1. Mapping of concepts

The foundation of this framework lies in treating physiological reserves as financial assets. Clinical default occurs when a patient’s health assets fall below their “debt,” defined as the metabolic requirements necessary to sustain life. Table 1 provides standardized operational definitions of variables used in the adapted Merton structural model for survival analysis.

**Table 1:** Mapping of clinical variables to parameters of the Merton jump–diffusion model.

| Variable category | Clinical variable | Operational definition (Model-Based) |
| --- | --- | --- |
| State variable | Body mass index (BMI) | Primary state variable $V$ representing the patient's physiological reserve. Modeled as a continuous stochastic process whose trajectory determines survival |
| Failure threshold | $K$ (Critical BMI threshold) | Default barrier defining biological failure. Death is assumed to occur when $V$ crosses below $K$ representing severe undernutrition |
| Time scale | $T$ (Exposure horizon) | Fixed follow-up window over which the probability of barrier crossing is estimated, applied consistently across all participants |
| Drift component | Age | Regression coefficient $\beta_{age}$ quantifying the effect of age on the drift of $V$, represents age-related decline in physiological recovery capacity |
| Drift component | Tuberculosis clinical form | Category-specific drift modifiers $\gamma_{TB}$ capturing differences in disease severity and progression across TB phenotypes (smear-positive, smear-negative, extrapulmonary) |
| Volatility component | HIV status | The volatility scaling factor $\phi_{HIV}$, applied to the baseline volatility $\sigma_0$, represents the increased health instability associated with immunosuppression |
| Jump intensity | Hospitalization status | Jump intensity modifier $\psi_{Hosp}$, which scales the frequency of acute health shocks ($\lambda$), with patients who are initially hospitalized assumed to experience more frequent destabilizing events |
| Jump magnitude | $\delta$ (Shock severity) | Standard deviation of jump sizes, representing the magnitude of sudden adverse health events such as opportunistic infections or clinical decompensation |

#### 2.5.2. Structural framework

The structural framework, or “health-balance sheet,” conceptualizes a patient’s physiological state as a dynamic system analogous to a company’s financial health. The model is built around three main pillars that determine the Distance-to-Death (DtD), equivalent to the distance-to-default, a measure representing the patient’s probability of survival. Fig 1 illustrates the interconnection between the three pillars. Pillar A includes asset dynamics and accounts for the trajectory of BMI, age, and TB classification. BMI is assumed to follow a stochastic process modeled by geometric Brownian motion, while age acts as a depreciation factor gradually reducing the patient’s recovery capacity, and TB classification functions like different industry sectors, each carrying its own baseline risk. Pillar B represents volatility and risk exposure factors. Certain factors can make a patient’s health trajectory less predictable, just as financial risk increases the volatility of a company’s assets. HIV co-infection is expected to amplify this volatility, causing greater fluctuations in health, while initial hospitalization increases the likelihood of sudden events or “clinical jumps,” representing an unstable environment. Pillar C governs the solvency indicator measured by the distance-to-default or Distance-to-Death (DtD). The DtD measures the distance to failure in terms of standard deviations from the default barrier. A high DtD indicates a patient with a strong health reserve, comparable to an investment-grade firm with low default risk, whereas a low DtD signals a patient in distress, at high risk of imminent mortality.

**Figure 1:**
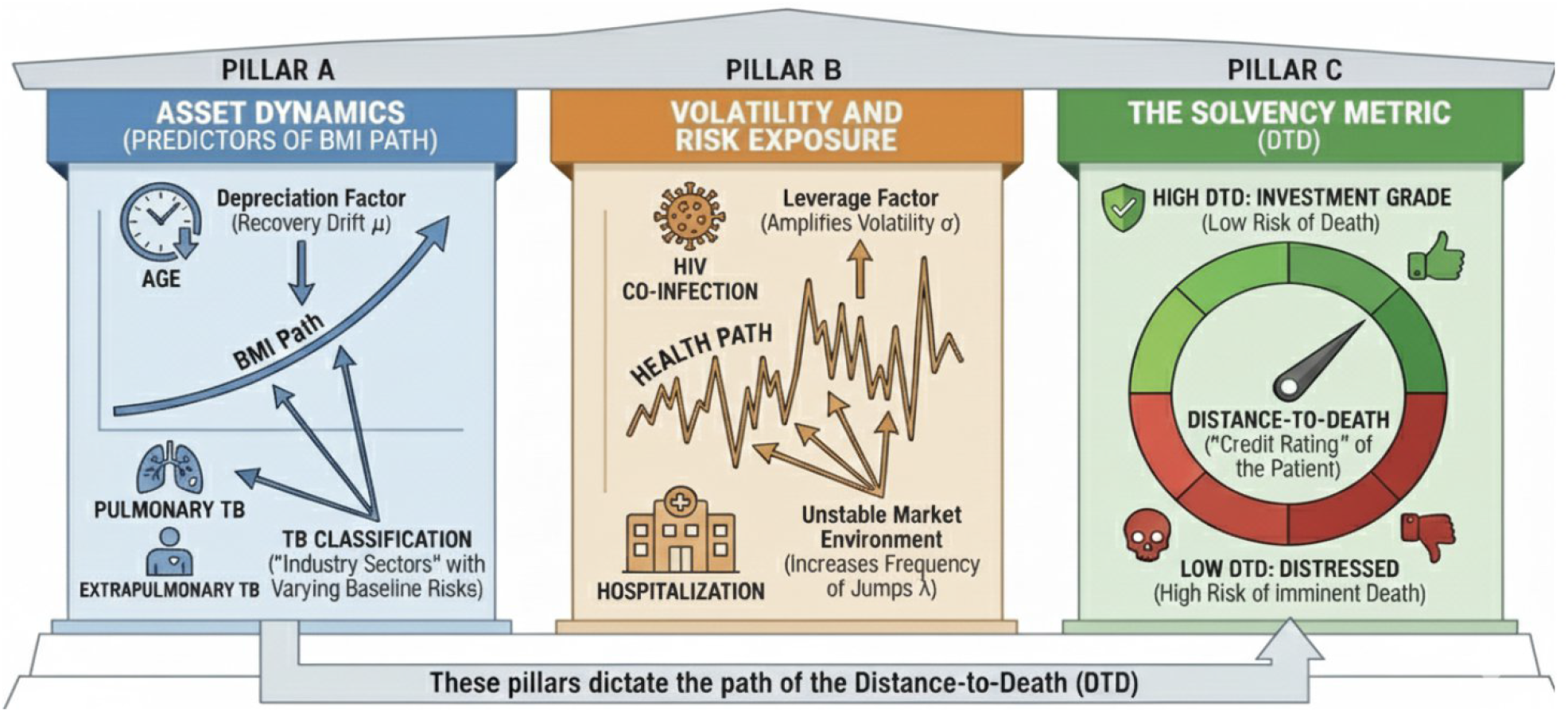
Structural framework.

#### 2.5.3. Conceptual equation flow

For each patient *i*, we denote the vector of clinical variables at admission as:

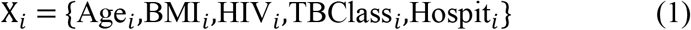

These variables are defined as follows: age at admission (*Age*_*i*_); body mass index at admission (*BMI*_*i*_); HIV status (*HIV*_*i*_ = 1 if positive, 0 if negative); tuberculosis classification (*TBClass*_*i*_ e.g. sputum-positive, sputum-negative, or extrapulmonary); and hospitalization status (*Hospit*_*i*_ = 1 if hospitalized, 0 otherwise). Sex and TB type (new vs. retreatment cases) were excluded from the model because they were not significantly associated with mortality.The mapping from clinical variables to the standardized risk score relies on a three-layer mathematical architecture. In this structural framework, survival is conceptualized as a state of biological solvency, whereby a patient remains alive only as long as their physiological reserve *V*_*T*_ stays above the critical threshold *K*. Mathematically, this condition *P*(*V*_*T*_ > *K*)is operationalized by standardizing the distance between the expected health trajectory and the failure barrier into a predictive survival probability.

##### 2.5.3.1. Structural layer: the stochastic differential equation (SDE)

The evolution of health capital (*V*_*t*_, measured by BMI) is considered to be governed by a jump-diffusion process. The fundamental equation is:

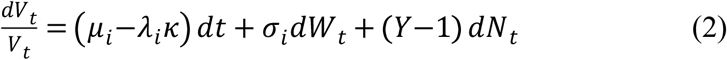

Where the diffusion component (*μ*_*i*_−*λ*_*i*_*κ dt* + *σ*_*i*_*dW*_*t*_) represents the continuous evolution of health with *μ*_*i*_ denoting the metabolic drift and *σ*_*i*_ the volatility or the physiological instability. The jump component (*Y*−1 *dN*_*t*_) captures sudden health shocks and *dN*_*t*_ is a Poisson process with intensity *λ*_*i*_, *Y*−1 represents the proportional loss of health capital during an acute event or a complication. The term *κ* = *E Y*−1 adjusts the drift to account for the expected effect of jumps, ensuring consistency of the overall process.

Applying the generalized Itô’s lemma for jump-diffusion processes, the analytical solution for the physiological reserve *V*_*t*_ at time *t* is given by:

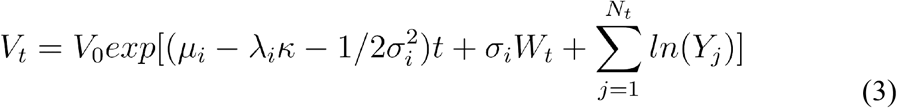

This expression shows that the patient’s health trajectory results from the interaction between a deterministic recovery trend, continuous metabolic fluctuations (*W*_*t*_), and the cumulative impact of discrete clinical shocks.

The parameter *κ* represents the expected relative change in the physiological reserve *V*_*t*_ when a clinical jump occurs. In accordance with the Merton framework, assuming that the magnitude of clinical shocks follows a log-normal distribution with a mean log-jump size of zero, *κ* is mathematically determined by the jump magnitude *δ* as follows:

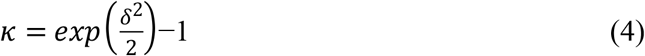

This relationship ensures that the drift adjustment in Equation (2) is internally consistent with the variability and severity of the acute clinical shocks captured by *δ*.

##### 2.5.3.2. Clinical parameter layer (Input layer)

The parameters of the stochastic differential equation were modeled as patient-specific functions of baseline characteristics. In particular, the individualized drift term was defined as:

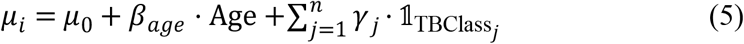

where *μ*_*i*_ represents the personalized metabolic drift of patient *i*, derived from a baseline drift *μ*_0_, adjusted for age through the coefficient *β*_*age*_, and further modified according to the TB clinical form. The term 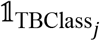 is an indicator variable equal to 1 if patient *i* belongs to TB class *j*, and 0 otherwise. Multiplying by *γ*_*j*_ adds the effect of that TB class to the personalized drift only if the patient is in that class.

The individualized volatility (physiological instability) was specified as:

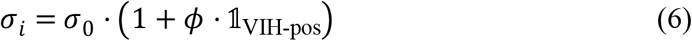

*σ*_*i*_ measures the physiological instability of patient *i* with *σ*_0_ the baseline volatility; The volatility increases if the patient is HIV positive (indicated by 𝟙_VIH-pos_) scaled by *ϕ*.

The individualized jump intensity was specified as:

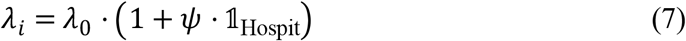

with *λ*_*i*_ quantifies the intensity of sudden health shocks; the base jump intensity *λ*_0_ is increased if the patient is hospitalized (indicated by 𝟙_Hospit_) scaled by *ψ*.

##### 2.5.3.3. Output layer: the Distance-to-Death (DTD) score

The DTD score is the analytical solution transforming the complex trajectory into a standardized measure (Z-score). Using the properties of the normal distribution and Merton approximation, we can state that:

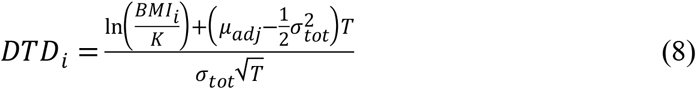

The aggregated parameters of the process were defined as follows:

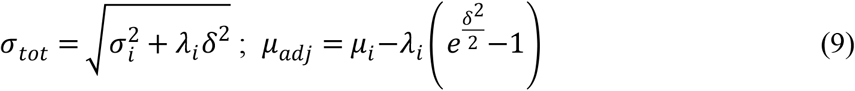

where *σ*_*tot*_ denotes the total volatility, integrating both continuous fluctuations and jump-related variability, and *μ*_*adj*_ is the drift adjusted for the expected impact of jumps. The Distance-to-Death score for patient *i* denoted *DtD*_*i*_, is defined relative to a critical threshold *K*, representing the level of physiological reserve (BMI) at which death occurs. In this framework, *δ* represents the average relative magnitude of jumps in health capital during acute complications, and *T* denotes the time horizon over which risk is evaluated.

#### 2.5.4. Benchmark model: the extended Cox proportional hazards model

To evaluate the added value of the structural approach, a frequentist survival model was developed as a comparator. While the standard Cox model assumes that the effect of covariates is constant over time, clinical reality in TB treatment often violates this “proportional hazards” assumption. We implemented an Extended Cox Model to account for non-proportionality by incorporating time-dependent effects (tt)[22]. The hazard function for patient *I* is defined as:

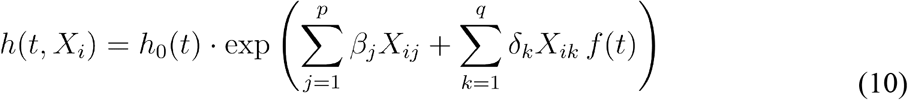

where h(t, X_*i*_) denotes the hazard function for patient *i*; *h*_0_ *t* is the baseline hazard; *X*_*ij*_ represents the value of the j^*th*^ fixed-effect covariate for patient *i*; *X*_*ik*_ denotes value of the k^*th*^ time-varying covariate for patient *i*; *β*_*j*_ are the coefficients associated with fixed effects; *δ*_*k*_ are the the coefficients associated with time-varying effects and *f(t)*=ln(t+1) is the time-transformation function.

#### 2.5.5. Statistical analysis

Data analysis was performed using R software version 4.5.2. The dataset was randomly split into a training set (70%), which was used to estimate and optimize model parameters, and a testing set (30%), which was reserved for evaluating the final predictive performance of the models.

##### 2.5.5.1. Implementation of the Merton model

The Merton jump-diffusion model was implemented as a structural survival framework to estimate the probability of “biological default” defined as death over a fixed time horizon corresponding to the maximum survival time in this cohort (240 days). The Merton model parameters were estimated via Maximum Likelihood Estimation (MLE) by minimizing the Negative Log-Likelihood (NLL) function. The Distance-to-Death (DTD) score is linked to the probability of survival by the equation:

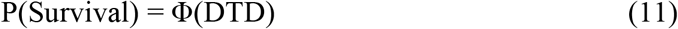

The NLL was calculated as follows:

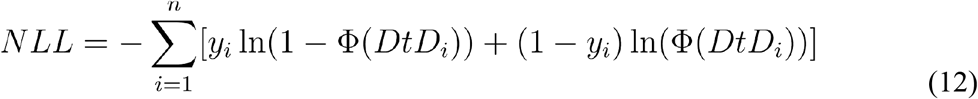

Here *n* is the total number of patients; *y*_*i*_ denotes the observed outcome for patient *i* (1 for death, 0 for survival); *ϕ*(*DtD*_*i*_) is the predicted probability of survival derived from the cumulative distribution function of the standard normal distribution applied to the DtD score; and 1−*ϕ*(*DtD*_*i*_) represents the predicted probability of death.

The optimization was performed using the L-BFGS-B algorithm, which allows for parameter estimation under box constraints. The following clinical parametrization was applied:

- Drift (μ_i_): modeled as a function of raw age and TB classification, with the “smear-positive pulmonary” group serving as the reference;
- Volatility (σ_i_) and Jump Intensity (λ_i_): adjusted via multiplicative coefficients for HIV-positive status and hospitalized patients, respectively;
- Critical Threshold (K): estimated within a biologically plausible range (15–30 kg/m^2^).

To evaluate the structural necessity of incorporating sudden health shocks, the full jump-diffusion model was compared with a nested pure-diffusion model, in which the jump intensity (λ) and jump magnitude (δ) were constrained to zero. Structural validation was performed using the likelihood ratio test (LRT) to assess whether the inclusion of jumps significantly improved the log-likelihood of the observed data. Model selection was further guided by the Akaike Information Criterion (AIC) to balance goodness-of-fit with model parsimony.

Standard errors for the Merton jump-diffusion model parameters were derived from the variance-covariance matrix obtained via the Moore-Penrose pseudo-inverse of the observed Hessian at the estimated parameters. Standard errors were computed as the square roots of the diagonal elements, ensuring non-negative values. From these, z-values and two-sided p-values were calculated assuming asymptotic normality, and results were summarized in a table reporting parameter estimates, standard errors, z-values, and p-values.

##### 2.5.5.2. Implementation of the comparator model

The extended Cox Proportional Hazards Model was used as the reference benchmark and was fitted to the training data. The proportional hazards assumption was verified using the Schoenfeld residuals test. For variables whose effects changed over time, time-transform functions (tt) were applied. Consequently, these variables were modeled as interactions with log(t+1), allowing their hazard ratios to evolve dynamically over the survival period. For prediction and calibration at the 240-day horizon, the cumulative baseline hazard of the extended Cox model was estimated using the Breslow method, with time-varying effects fixed at t=240 to ensure comparability with the Merton model’s fixed-horizon estimates.

##### 2.5.5.3. Estimation of the extended Cox model

The coefficients β of the extended Cox model were optimized using the Newton-Raphson algorithm by maximizing the log-partial likelihood *ℓ(β)*, which is expressed as:

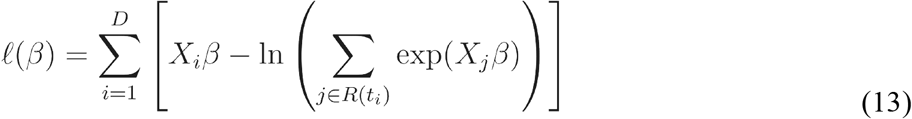

For each event time *t*_*i*_ the risk score *X*_*i*_*β* was computed as the linear combination of predictors for the patient who experienced the event, while the risk set *R*(*t*_*i*_) included all individuals still at risk alive and uncensored just prior to *t*_*i*_. This approach allowed us to account for both fixed and time-varying covariates, providing a flexible framework to model the instantaneous hazard of death during TB treatment.

##### 2.5.5.4. Calibration and model performance

The calibration and discriminative power of the models were evaluated in the test set. Graphical methods were employed to assess calibration by plotting deciles of predicted survival probabilities against observed Kaplan-Meier estimates. Discrimination was measured using Harrell’s C-index; for the Merton model, this was based on the negative Distance-to-Death (-DTD) score, while the linear predictor was used for the extended Cox model. To assess the statistical significance of performance differences, a non-parametric bootstrap procedure (1,000 iterations) was used to generate 95% confidence intervals (CI) and bootstrap-based p-values for the difference in C-indices.

## 3. Results

### 3.1. Study population

A total of 28,360 TB patients were registered, of whom 15,182 were included in the final analysis after excluding 13,178 individuals. The main reason for exclusion was unknown HIV status (n = 9,023; 68.5% of exclusions), followed by missing follow-up duration (n = 3,612), unspecified hospitalization status (n = 2,692), and data issues related to body weight (n = 33). Missing data were most frequent in the early 2000s, prior to the systematic integration of TB/HIV services and routine HIV testing in Cameroon, indicating that the included cohort reflects a period with improved diagnostic completeness. A comparative analysis of baseline characteristics between included and excluded participants revealed statistically significant differences across all evaluated parameters, likely driven by the large sample size and resulting statistical power. Excluded patients were more likely to be managed as outpatients (19.5% vs. 14.7%) and presented with higher proportions of EPTB (25.8% vs. 13.7%) and SNPTB (19.5% vs. 8.2%) compared to the included cohort. Differences in sex distribution (58.1% male in the included group vs. 59.9% in the excluded group) and patient category (91.5% new cases in the included group vs. 92.7% in the excluded group) were statistically significant but clinically minor.

The study population was predominantly male (58.1%) with a median age of 33 years [interquartile range (IQR) : 26–43]. At admission, the median BMI was 20.7 kg/m ^2^ (IQR: 18.3–23.0). HIV co-infection was present in 35.4% of the cohort, and 85.3% of patients were initialy hospitalizated. Regarding TB classification, 78.1% had smear-positive pulmonary TB (SPPTB), 8.2% had smear-negative pulmonary TB (SNPTB), and 13.7% had extrapulmonary TB (EPTB). The overall mortality rate during the follow-up period was 7.0% (n = 1,059).The median time to death was 44.2 days (IQR: 26–59), with 55.1% of deaths occurring within the first 30 days. The general characteristics of the study participants are summarized in Table 1.

**Table 1:** General characteristics of included patients.

| Characteristics | N = 15,182 (%) |
| --- | --- |
| Sex |  |
| Male | 8,816 (58.1) |
| Female | 6,366 (41.9) |
| Age, yrs, median (25<sup>th</sup> - 75<sup>th</sup> percentiles) | 33 (26 - 43) |
| Type patient |  |
| New cases | 13,892 (91.5) |
| Retreatment cases | 1,290 (8.5) |
| TB location |  |
| SPPTB | 11,853 (78.1) |
| SNPTB | 1,248 (8.2) |
| EPTB | 2,081 (13.7) |
| HIV status |  |
| Negative | 9,809 (64.6) |
| Positive | 5,373 (35.4) |
| BMI, kg/m<sup>2</sup> |  |
| Median (25<sup>th</sup> - 75<sup>th</sup> percentiles) | 20.7 (18.3 - 23.0) |
| < 18.5 kg/m<sup>2</sup> |  |
| Initial hospitalization |  |
| Yes | 12,953 (85.3%) |
| No | 2,229.0 (14.7%) |
| TB outcome |  |
| Favorable | 11,834 (77.9) |
| Failure | 89 (0.6) |
| Loss to follow-up | 2,200 (14.5) |
| Death | 1,059 (7.0) |

HIV, human immunodeficiency virus TB, tuberculosis ; SPPTB, smear positive pulmonary TB ; SNPTB, smear negative pulmonary TB ; EPTB, extrapulmonary TB ; BMI, body mass index

### 3.2. Benchmark Model: Extended Cox proportional hazards

The multivariable frequentist survival analysis identified several baseline factors significantly associated with mortality. Lower BMI at admission was independently associated with an increased hazard of death (hazard ratio [HR] per unit increase: 0.88; 95% confidence interval [CI]: 0.86–0.90; p < 0.001). HIV-positive status (HR: 5.31; 95% CI: 3.48–8.09; p < 0.001) and initial hospitalization (HR: 11.28; 95% CI: 3.84–33.16; p < 0.001) were associated with the highest mortality risks. Both SNPTB (HR: 2.66; 95% CI: 2.18–3.25; p < 0.001) and EPTB) (HR: 2.60; 95% CI: 2.20–3.09; p < 0.001) were associated with significantly higher hazards compared with the smear-positive pulmonary TB reference group.Time-dependent effects indicated that the hazards associated with HIV infection and hospitalization decreased over the course of treatment (both p < 0.01), suggesting a time-varying impact of these factors on mortality risk (Table 2).

**Table 2:** Extended Cox model results.

| Parameter | Coefficient ($\beta$) | Hazard Ratio (95% CI) | p-value |
| --- | --- | --- | --- |
| Age, yrs (per unit increase) | 0.0245 | 1.025 (1.019–1.030) | <0.001 |
| BMI, kg/m<sup>2</sup> ( per unit increase) | -0.1216 | 0.885 (0.867–0.904) | <0.001 |
| HIV positive | 1.6698 | 5.311 (3.484–8.096) | <0.001 |
| Hospitalized participants | 2.4238 | 11.288 (3.842–33.167) | <0.001 |
| TB class |  |  |  |
| SPPTB | Reference | Reference | / |
| SNPTB | 0.9584 | 2.661(2.175–3.254) | <0.001 |
| EPTB | 0.9584 | 2.608 (2.199–3.092) | <0.001 |
| HIV positive (time-varying) | -0.1261 | 0.881 (0.782–0.994) | 0.040 |
| Hospitalized (time-varying) | -0.4377 | 0.646 (0.488–0.853) | 0.002 |

BMI, body mass index ; CI, confidence interval, HIV, human immunodeficiency virus TB, tuberculosis ; SPPTB, smear positive pulmonary TB ; SNPTB, smear negative pulmonary TB ; EPTB, extrapulmonary TB

### 3.3. Structural Merton Jump-Diffusion parameters

The estimated parameters of the Merton jump-diffusion model are presented in Table 3.The structural Merton model successfully converged, identifying a critical nutritional threshold (K, BMI) of 17.329 kg/m^2^ (standard error [SE]: 1.006; p < 0.001), representing the physiological barrier for biological failure.The dynamics of health capital were significantly influenced by key clinical covariates. The metabolic drift (μ), reflecting the continuous trajectory of recovery or decline in physiological reserve, was estimated at a baseline value of 3.089. This drift decreased significantly with increasing age (β_age_ = −0.017; p < 0.001), indicating a faster depletion of physiological reserves in older patients. TB clinical forms also exerted a significant negative effect on drift. Compared with SPPTB, patients with SNPTB and EPTB showed reduced recovery trends of −0.657 and −0.647, respectively (both p < 0.001).

**Table 3:** Parameters of the Merton jump-diffusion model.

| Parameter | Estimate | SE | z-value | p-value |
| --- | --- | --- | --- | --- |
| $\mu_0$ (baseline drift) | 3.090 | 0.282 | 10.96 | <0.001 |
| $\beta_{age}$ (age effect on drift) | -0.017 | 0.001 | -11.48 | <0.001 |
| $\sigma_0$ (baseline volatility) | 0.737 | 0.011 | 65.58 | <0.001 |
| $\lambda_0$ (baseline jump intensity) | 0.326 | 0.490 | 0.67 | 0.505 |
| K (critical BMI threshold) | 17.329 | 1.006 | 17.23 | <0.001 |
| $\delta$ (jump magnitude) | 0.435 | 0.365 | 1.19 | 0.233 |
| $\gamma_1$ (TB class-EPTB) | -0.647 | 0.059 | -10.97 | <0.001 |
| $\gamma_2$ (TB class-SNPTB) | -0.657 | 0.074 | -8.84 | <0.001 |
| $\phi$ (HIV effect on volatility) | 1.446 | 0.023 | 62.64 | <0.001 |
| $\psi$ (hospitalization effect on jumps) | 4.263 | 3.918 | 1.09 | 0.277 |

The baseline physiological volatility (σ), reflecting underlying metabolic instability, was estimated at 0.737. HIV co-infection emerged as a major destabilizing factor, significantly increasing volatility by a multiplicative factor of 1.446 (ϕ_HIV_; p < 0.001). Regarding acute health shocks, the baseline jump intensity (λ) was 0.326. Although initial hospitalization was associated with an increased frequency of these shocks (multiplier: 4.263), this effect did not reach statistical significance (p = 0.277), suggesting that its impact may be partially mediated by other covariates. However, model comparison strongly supported the inclusion of the jump component, as the full jump-diffusion model provided a significantly better fit than the pure diffusion model (ΔAIC = −17.49; LRT χ^2^ = 23.48, p < 0.001), indicating that acute health shocks contribute meaningfully to mortality risk beyond individual predictors.

### 3.4. Model performance and comparison

In the independent test set, the Merton Jump-Diffusion model demonstrated slightly superior discriminative capacity relative to the Extended Cox benchmark. The structural framework achieved a Harrell’s C-index of 0.781 (95% CI: 0.756–0.807), slightly outperforming the Extended Cox model, which achieved a C-index of 0.772 (95% CI: 0.745–0.798). This absolute improvement of 0.009 in the C-index was statistically significant (bootstrap p = 0.038), confirming the enhanced predictive accuracy of the structural approach in this clinical context. The comparative calibration of the structural Merton Jump-Diffusion model and the Extended Cox benchmark across risk deciles is illustrated in Fig 2. Visual inspection of the 240-day calibration plots revealed that the Merton Jump-Diffusion model achieved higher precision in elevated-risk deciles, following the ideal diagonal more closely than the Extended Cox model, which deviated at higher mortality probabilities.

**Fig 2.**
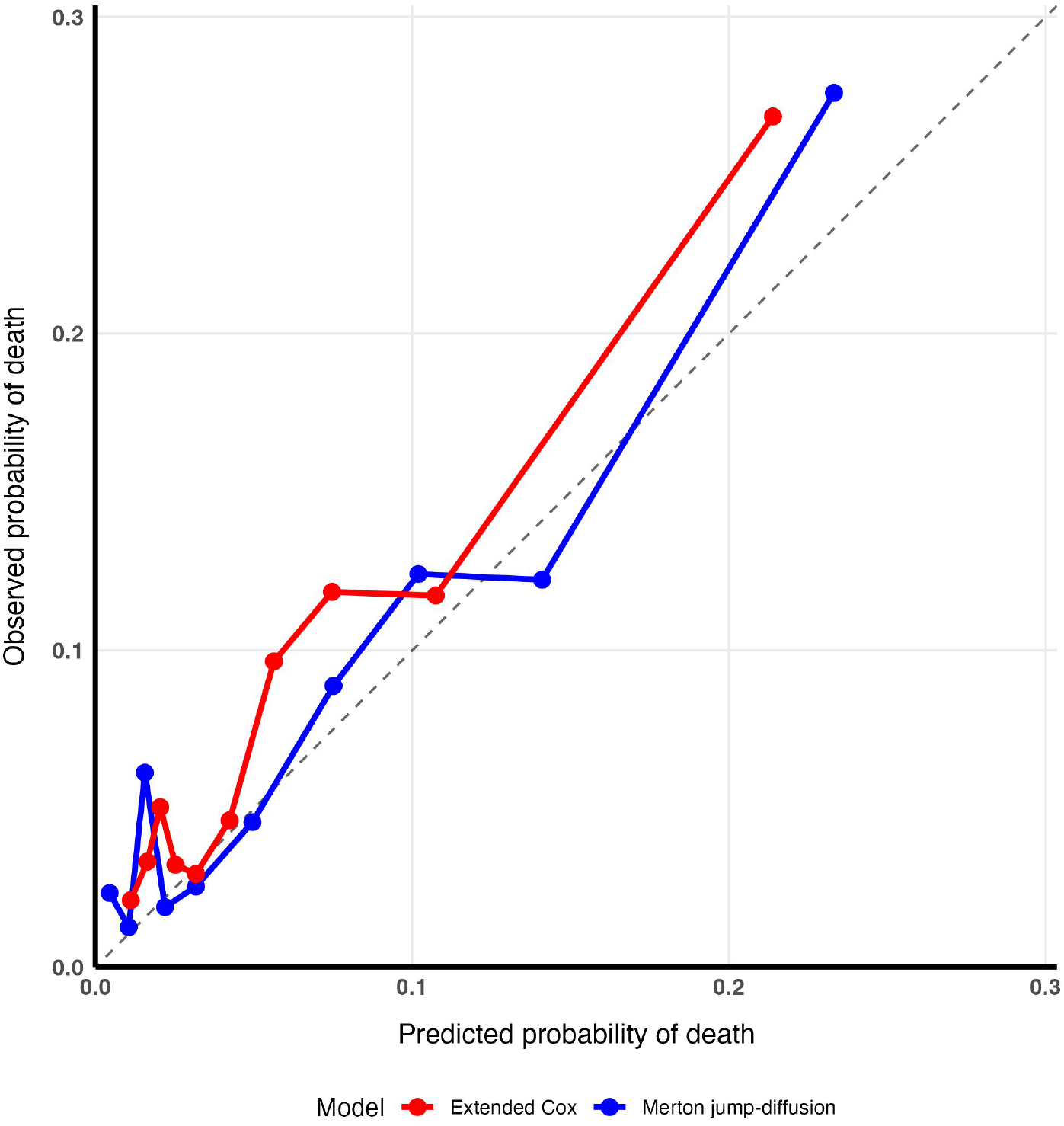
Comparative calibration of structural and associative models for tuberculosis mortality risk.

### 3.5. Distribution and clinical relevance of the DtD score

Fig 3 presents the validation and risk stratification performance of the structural DtD score, illustrating its ability to discriminate survival outcomes, capture individualized risk patterns, and stratify patients into clinically meaningful risk groups.The Distance-to-Death (DtD) score stratified mortality risk in the test cohort (N = 4,555). Survivors (n = 4,241) had a median DtD of 1.83 (IQR: 1.33–2.16), whereas non-survivors (n = 314) had a median DtD of 1.12 (IQR: 0.74–1.46). Admission BMI correlated positively with DtD. Patients with identical BMI values exhibited a range of DtD scores, varying with age, HIV co-infection status, and TB phenotype. Kaplan-Meier analysis across DtD quartiles revealed a significant, non-linear mortality gradient (log-rank p < 0.0001). Mortality was highest in the first quartile (Q1: DtD <1.28) at 16.7% with a median DtD of 1.00. Mortality decreased to 7.5% in Q2 (DtD 1.28–1.78; median DtD 1.52), 1.8% in Q3 (DtD 1.78–2.14; median DtD 1.97), and 1.6% in Q4 (DtD > 2.14; median DtD 2.34).

**Fig 3.**
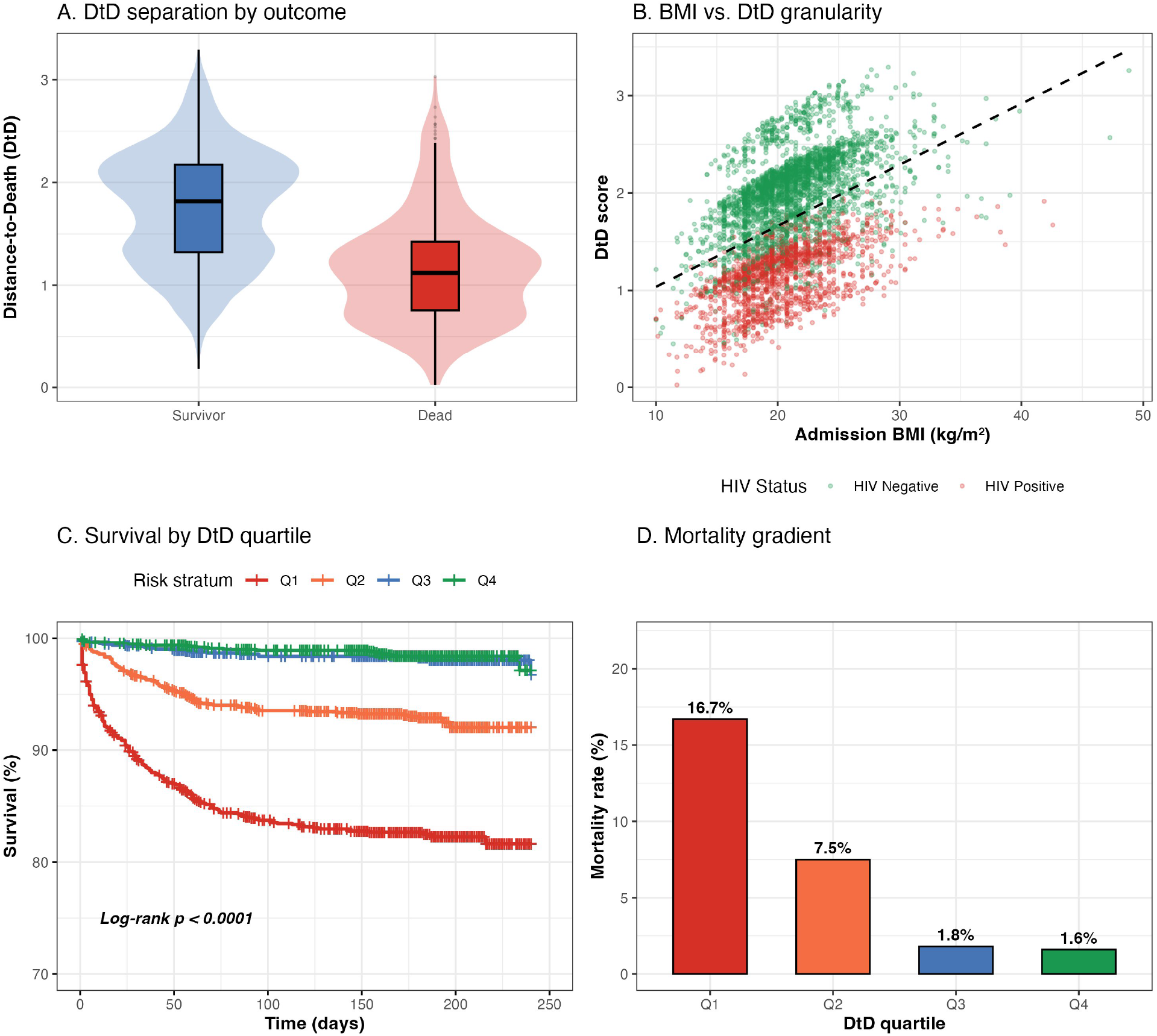
Validation and risk stratification of the structural Distance-to-Death (DtD) score in tuberculosis. (A) Baseline DtD distribution stratified by survival outcome; (B) Body mass index (BMI) vs. DtD granularity demonstrating individualized risk assessment by HIV status; (C) Kaplan-Meier survival analysis across high-contrast DtD risk quartiles; (D) Observed mortality rates within the four risk strata in the testing set.

### 3.6. Clinical implementation: The DtD-TB Decision-Support tool

To translate the structural Merton jump-diffusion framework into clinical practice, we developed an interactive DtD-TB digital tool that calculates a patient’s Distance-to-Death (DtD) score at admission using age, BMI, HIV status, TB form, and initial management mode. The tool computes three prognostic indicators: the DtD Z-score relative to the critical survival threshold (K = 17.329 kg/m^2^), survival probability over 240 days (Φ(DtD)), and exact death risk (P(Death) = Φ(−DtD)). Patients are automatically stratified into four risk levels based on probability of death: Critical (>10% risk) for imminent biological insolvency requiring intensive monitoring and nutritional support; High (5–10% risk) requiring closer follow-up; Moderate (2.5–5% risk) indicating relative stability; and Low/Stable (<2.5% risk) with strong biological solvency and minimal early mortality risk. The DtD-TB decision-support tool is available as an interactive online application for clinical use at: [DtD-TB tool risk assessent](https://structural-jump-diffusion-fiimxadjs8uwgtn4skzjh8.streamlit.app/).

## 4. Discussion

To the best of our knowledge, this study represents one of the first applications of a structural jump-diffusion modeling framework, inspired by quantitative finance, to the analysis of time-to-event outcomes in biomedical research and clinical epidemiology. By introducing the concept of Distance-to-Death (DtD), we propose a shift from purely associative approaches, such as the Cox model, toward a mechanistic perspective in which survival is driven by the dynamic evolution of a physiological reserve relative to a critical biological threshold. This framework is not restricted to TB but may be generalized to a wide range of biomedical conditions characterized by progressive decline and acute clinical events.

The main finding of this study is the modest but statistically significant improvement in predictive performance of the DtD model compared with the extended Cox model (C-index: 0.781 vs 0.772; p = 0.038). Although this improvement is modest in magnitude, the primary value of the structural approach lies in its complementary nature, as it offers mechanistic interpretability through the DtD score, a feature not captured by traditional associative models. While the Cox model estimates the average effect of covariates on the instantaneous hazard, the Merton framework explicitly models the stochastic trajectory of the biomarker. This result is consistent with the ‘process point of view’ advocated by Aalen and Gjessing[17], whereby survival is conceptualized as a first-passage time to an absorbing state, providing a mechanistic interpretation of hazard dynamics that is not captured by traditional associative models.

The model identified a critical BMI threshold (K) of 17.329 kg/m^2^, which is consistent with established criteria for severe malnutrition, typically defined as a BMI below 17.8 kg/m^2^[23]. As this threshold is approached, biological solvency becomes progressively compromised. Prior studies have highlighted that BMI serves as a proxy for underlying metabolic and nutritional reserve capacity, which is closely linked to survival outcomes [6,24,25]. The DtD metric extends this concept by quantifying the margin of safety relative to this critical boundary at admission.

Within the structural framework, the drift (μ) represents the patient’s tendency toward recovery or deterioration. Age had a significant negative effect on drift (β_age_ = −0.017, p < 0.001), reflecting a progressive decline in physiological resilience, consistent with immunosenescence and reduced metabolic plasticity that limit the ability of older individuals to rebuild body reserves during infection[26–28]. In addition, SNPTB and EPTB exhibited markedly more unfavorable drift values compared with SPPTB (−0.647 and −0.657, respectively). These forms are associated with delayed diagnosis or underlying immunosuppression in SNPTB and systemic dissemination in EPTB[29,30], contributing to a more complex catabolic burden and less predictable response to treatment.

Another key contribution of the model is the parameterization of physiological instability through volatility (σ). HIV infection increased volatility by a factor of 1.446 (p < 0.001), indicating more unpredictable health trajectories in HIV-TB co-infected patients. Within the Merton framework, this translates into a higher probability of crossing the failure threshold. Clinically, this instability is consistent with the well-documented susceptibility of HIV-infected individuals to opportunistic infections and immune reconstitution inflammatory syndrome (IRIS), both of which can lead to abrupt clinical deterioration despite apparently stable baseline status[5,31,32].

Incorporating jump components proved statistically necessary (*p*<0.001, LRT), indicating that TB-related mortality cannot be explained solely by gradual decline. Instead, it is punctuated by acute clinical shocks such as haemoptysis, septic shock, or sudden decompensation. By assuming a mean log-jump size of zero, we modeled these complications as sources of pure stochastic uncertainty rather than systematic trends. In this structural configuration, the parameter *κ* representing the expected relative change in physiological reserve during a shock is determined entirely by the estimated jump magnitude (*δ*). This allowed the framework to clearly distinguish between the predictable, progressive erosion of health reserves (captured by the personalized drift) and the unpredictable jumps that abruptly narrow the patient’s distance to biological insolvency. Although the hospitalization-related multiplier did not reach individual statistical significance (p = 0.277), the superiority of the full model over the pure diffusion model confirms that the risk structure is inherently discontinuous. This observation aligns with findings by Zachariah et al. on the rapid onset of early mortality in resource-limited settings[33].

The DtD score provides a practical framework for clinical stratification. Approximately 60% of deaths occurred in the lowest quartile (DtD < 1.28), identifying a subgroup with immediate biological vulnerability. In contrast, patients with DtD > 2.14 had a very low mortality rate (1.6%). This stratification may inform prioritization of intensive nutritional support or closer clinical monitoring.

The DtD-TB tool translates the structural Merton Jump-Diffusion model into clinical practice, providing an objective assessment of patient risk at admission using five standard parameters. By integrating physiological reserves and key modifiers such as age, HIV status, and TB phenotype, it enables mechanistic mortality predictions and automatic stratification into four actionable risk levels. This facilitates timely triage, targeted interventions, and efficient resource allocation in high-burden settings. Its online, interactive format supports rapid adoption and standardized risk assessment, demonstrating the feasibility of applying structural models from quantitative finance to clinical epidemiology. Future prospective studies are warranted to validate the tool’s predictive performance across diverse populations and care settings.

Several limitations of this study warrant consideration. First, the retrospective nature of the 20-year study period encompasses substantial changes in tuberculosis and HIV management, including the rollout of antiretroviral therapy and improvements in diagnostic practices. Although the model is designed to capture underlying biological dynamics, such temporal heterogeneity may have influenced the stability of the estimated parameters across different eras. Second, body mass index (BMI) was calculated using standardized height values by sex, a pragmatic approach that enables consistent computation across a large retrospective dataset where individual height measurements were unavailable or incomplete. This strategy ensures comparability of BMI values while maintaining analytical feasibility, and is appropriate for large-scale structural modeling where relative differences in body mass are of primary interest. Third, the framework is mechanistically inspired, as it reconstructs a latent health trajectory without relying on dense longitudinal measurements of BMI. While BMI serves as a useful proxy for nutritional and metabolic reserve, it does not fully capture the multidimensional nature of patient severity. Future extensions of the health asset (V_t_) could incorporate additional clinical and laboratory markers, such as baseline bacillary load, inflammatory biomarkers (e.g., CRP), and functional status indicators (e.g., Karnofsky score). Finally, the model assumes a constant survival threshold (K), whereas the true clinical failure boundary is likely to vary across individuals depending on comorbidities, frailty, and other unobserved factors.

Despite these considerations, the large sample size, the consistent direction and magnitude of the estimated effects, and the robustness observed in out-of-sample validation support the stability of the findings. Moreover, the structural nature of the model grounded in relative dynamics rather than absolute values helps mitigate the impact of measurement imprecision and temporal heterogeneity, thereby supporting the overall validity of the results. Future refinements of this structural framework may include the empirical estimation of a non-zero mean log-jump component (α), allowing for systematic downward shifts in health capital associated with clinical complications. In addition, extending the model from a constant failure barrier (K) to a dynamic threshold that adapts to individual comorbidity profiles or age-related frailty may further improve the discriminative performance of the Distance-to-Death (DtD) metric.

## 5. Conclusion

The Merton jump-diffusion model introduces a structural, mechanistic perspective to survival modeling in biomedical contexts, specifically within the clinical epidemiology of tuberculosis. By incorporating the Distance-to-Death (DtD) metric, the model captures individual recovery dynamics, age-dependent effects, HIV-associated physiological instability, and acute clinical shocks. This framework moves beyond purely associative approaches, providing a patient-centered view of risk grounded in the trajectory of physiological resilience. While this study establishes the DtD score as a robust proof-of-concept for tuberculosis triage, subsequent iterations of the model should explore the integration of stochastic volatility and time-varying parameters to more accurately reflect the evolving physiological resilience of patients as they progress from the intensive to the continuation phase of treatment.

## Author contributions

E.W.P-Y conceived the study, designed the modeling framework, performed analyses, and drafted the manuscript; E.H.P-Y contributed to model development and critically revised the manuscript; H.L.N.P-Y curated and managed clinical data and assisted with manuscript preparation; A.D contributed to clinical data collection and ethical oversight; A.D.B facilitated data acquisition and provided clinical insights. All authors read and approved the final manuscript.

### Competing interests

The authors have declared that no competing interests exist.

### Compliance with ethical standards

This study was conducted in accordance with the principles of the Declaration of Helsinki. Ethical approval was obtained from the Faculty of Medicine and Biomedical Sciences of the University of Yaoundé 1 (No. 0195/2023).

### Financial disclosures

This research received no specific grant from any funding agency in the public, commercial, or not-for-profit sectors.

### Data availability and code availability

Data are available from the corresponding author upon reasonable request; however, due to ethical and privacy restrictions regarding patient clinical records, the original dataset is not publicly available. To ensure verification and reproducibility of the Merton Jump-Diffusion framework, a statistically representative synthetic dataset along with all computational code used for data processing, model development, and analysis are provided at https://github.com/pefura/structural-jump-diffusion.This implementation includes the full pipeline for the structural model, the computation of the Distance-to-Death (DtD) metric, and the scripts required for the replication of all results presented in this manuscript.

